# Machine learning on drug-specific data to predict small molecule teratogenicity

**DOI:** 10.1101/860627

**Authors:** Anup P. Challa, Andrew L. Beam, Min Shen, Tyler Peryea, Robert R. Lavieri, Ethan S. Lippmann, David M. Aronoff

## Abstract

Pregnant women are an especially vulnerable population, given the sensitivity of a developing fetus to chemical exposures. However, prescribing behavior for the gravid patient is guided on limited human data and conflicting cases of adverse outcomes due to the exclusion of pregnant populations from randomized, controlled trials. These factors increase risk for adverse drug outcomes and reduce quality of care for pregnant populations. Herein, we propose the application of artificial intelligence to systematically predict the teratogenicity of a prescriptible small molecule from information inherent to the drug. Using unsupervised and supervised machine learning, our model probes all small molecules with known structure and teratogenicity data published in research-amenable formats to identify patterns among structural, meta-structural, and *in vitro* bioactivity data for each drug and its teratogenicity score. With this workflow, we discovered three chemical functionalities that predispose a drug towards increased teratogenicity and two moieties with potentially protective effects. Our models predict three clinically-relevant classes of teratogenicity with AUC = 0.8 and nearly double the predictive accuracy of a blind control for the same task, suggesting successful modeling. We also present extensive barriers to translational research that restrict data-driven studies in pregnancy and therapeutically “orphan” pregnant populations. Collectively, this work represents a first-in-kind platform for the application of computing to study and predict teratogenicity.

## Introduction

### 1.1. Risky prescriptive behavior in pregnancy

*Teratogenicity* is the most serious manifestation of iatrogenic fetal toxicity: teratogens lead to fetal malformation and are implicated in lifelong physical and/or mental disabilities^1^. Nonetheless, clinical trial results of drug exposure during pregnancy are often conflicting^2–4^, and teratogenicity scoring for small molecules is unsystematic and performed outside the clinical environment^5–7^. The consequences of this subjectivity are seen in the high rate of unintended maternal exposure to a teratogenic agent^8^, reminiscent of the “thalidomide disaster” of the early 1960s^9,10^. Following this disaster, randomized, controlled trials (RCTs) were modified to exclude pregnant populations, fearing unintended teratogenicity from exposure to unsystematically profiled drugs^10^. This change continues to “orphan” pregnant women, as many diseases in women’s health lack safe and effective drug choices for treatment^8, 11, 12^.

In the wake of the “thalidomide disaster,” the United States Food and Drug Administration (FDA) developed a five-point scale for ranking the teratogenicity of a compound^7–9, 11^. This scale is presented in Table 1 (Appendix).

**Table 1.**
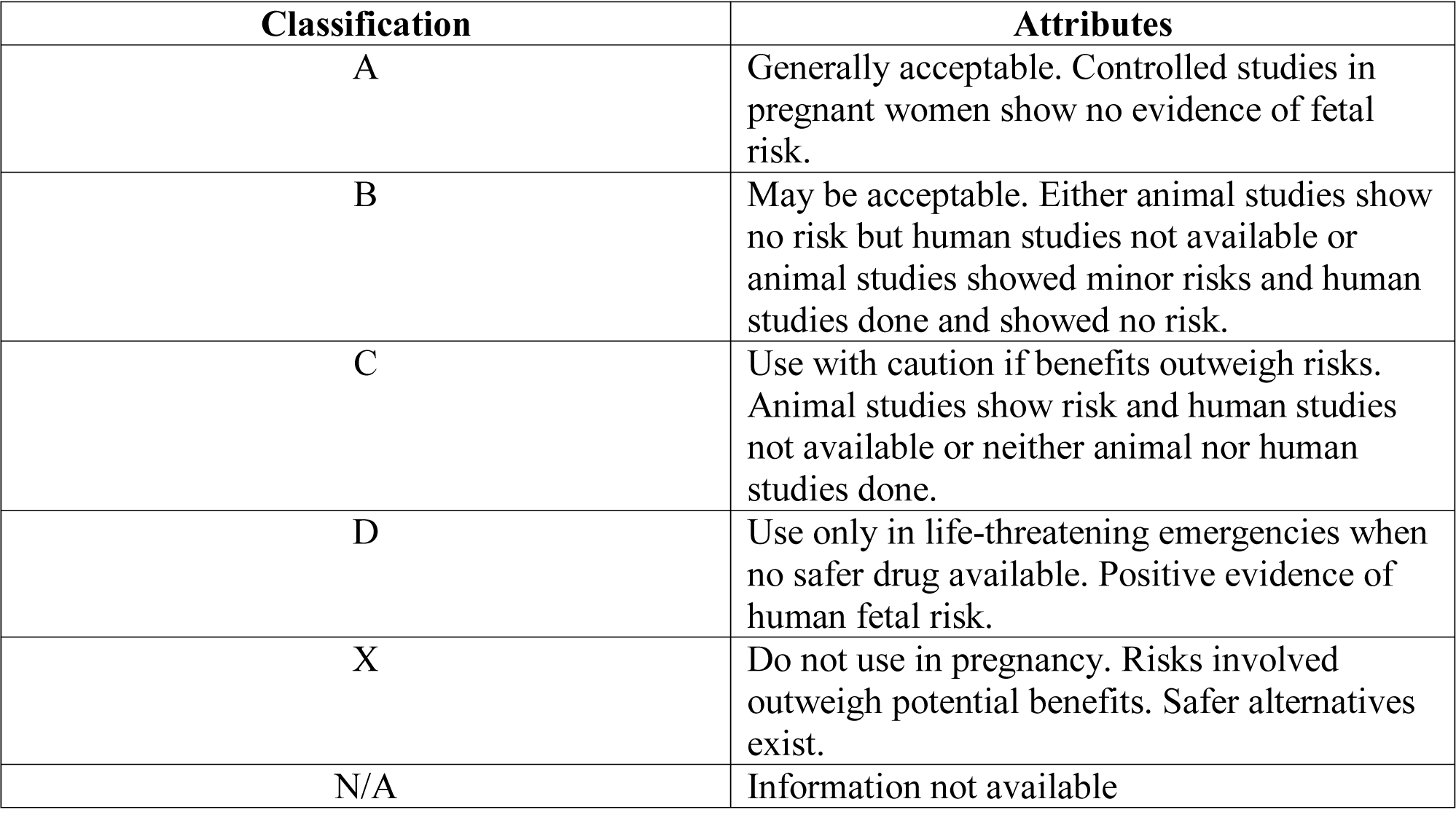
Teratogenicity scoring criteria established by the FDA are driven by a lack of human data, making them dangerously imprecise for application at the bedside^7,13^.

| Classification | Attributes |
| --- | --- |
| A | Generally acceptable. Controlled studies in pregnant women show no evidence of fetal risk. |
| B | May be acceptable. Either animal studies show no risk but human studies not available or animal studies showed minor risks and human studies done and showed no risk. |
| C | Use with caution if benefits outweigh risks. Animal studies show risk and human studies not available or neither animal nor human studies done. |
| D | Use only in life-threatening emergencies when no safer drug available. Positive evidence of human fetal risk. |
| X | Do not use in pregnancy. Risks involved outweigh potential benefits. Safer alternatives exist. |
| N/A | Information not available |

A hallmark of the binning within this scale is the absence of definitive human data: at present, teratogenicity scores are established pre-clinically by pharmacologists, who evaluate biomarkers of fetal toxicity in animal models^5,6^. This approach is inherently limited, as common *in vivo* models are not sufficiently representative of human physiology^13^, and human subjects are not included in the teratogenicity scoring process for ethical reasons^11, 14, 15^. Indeed, the limited human data available for teratology scoring are often derived retrospectively from high-profile cases of fetal malformation resulting from drug exposure^9, 16, 17^. While new FDA standards for scoring teratogenicity acknowledge these limitations by providing fewer, more holistic toxicity scores, these standards still suffer from the absence of robust human data and are not yet integrated in clinical decision-making tools^18^.

Collectively, the factors above create a significant degree of uncertainty at the point of care (POC), as providers are guided on contradictory, incomplete, and non-human derived information in their choice of prescriptions for pregnant women. This dilemma is of special consequence to expectant mothers with chronic morbidities pre-existing to their pregnancies^11^.

### 1.2. Target rationale for teratogenesis

Fetal exposure to a teratogen *in utero* strongly associates with cognitive and/or physical disabilities, resulting from dysregulation of key developmental processes such as neurulation, purine and pyrimidine synthesis, and lipid anabolism^2,19^.

Broadly, teratogens may be categorized by their mechanism of action (MOA) as either “on-target” or “off-target^20–22^.” “On-target” teratogenicity implies the generation of adverse phenotypes from bioactive agents impacting well-defined protein targets that are critically regulated in development. In contrast, “off-target” teratogenicity implies *mutagenicity*, resulting from DNA damage such as alkylation and thymine dimerization. “Off-target” teratogenicity involves repeated reactions between a teratogen and newly-synthesized nucleic acid residues, often resulting from the generation of reactive oxygen species (ROS) generated from drug metabolism^20^.

Thus, teratology is known to converge on few principal MOA classes^19,23^, which are outlined in Table 2.

**Table 2.**
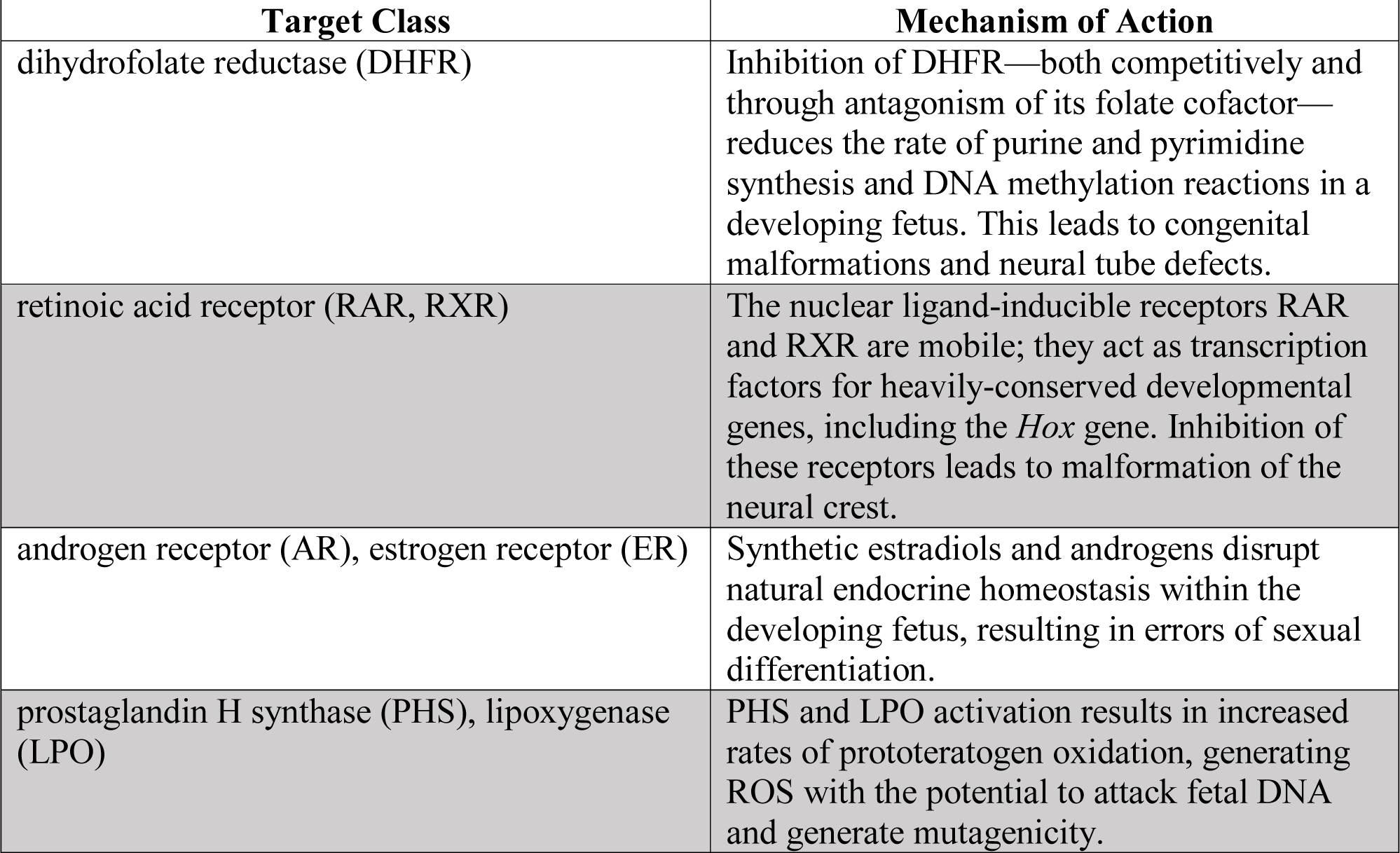
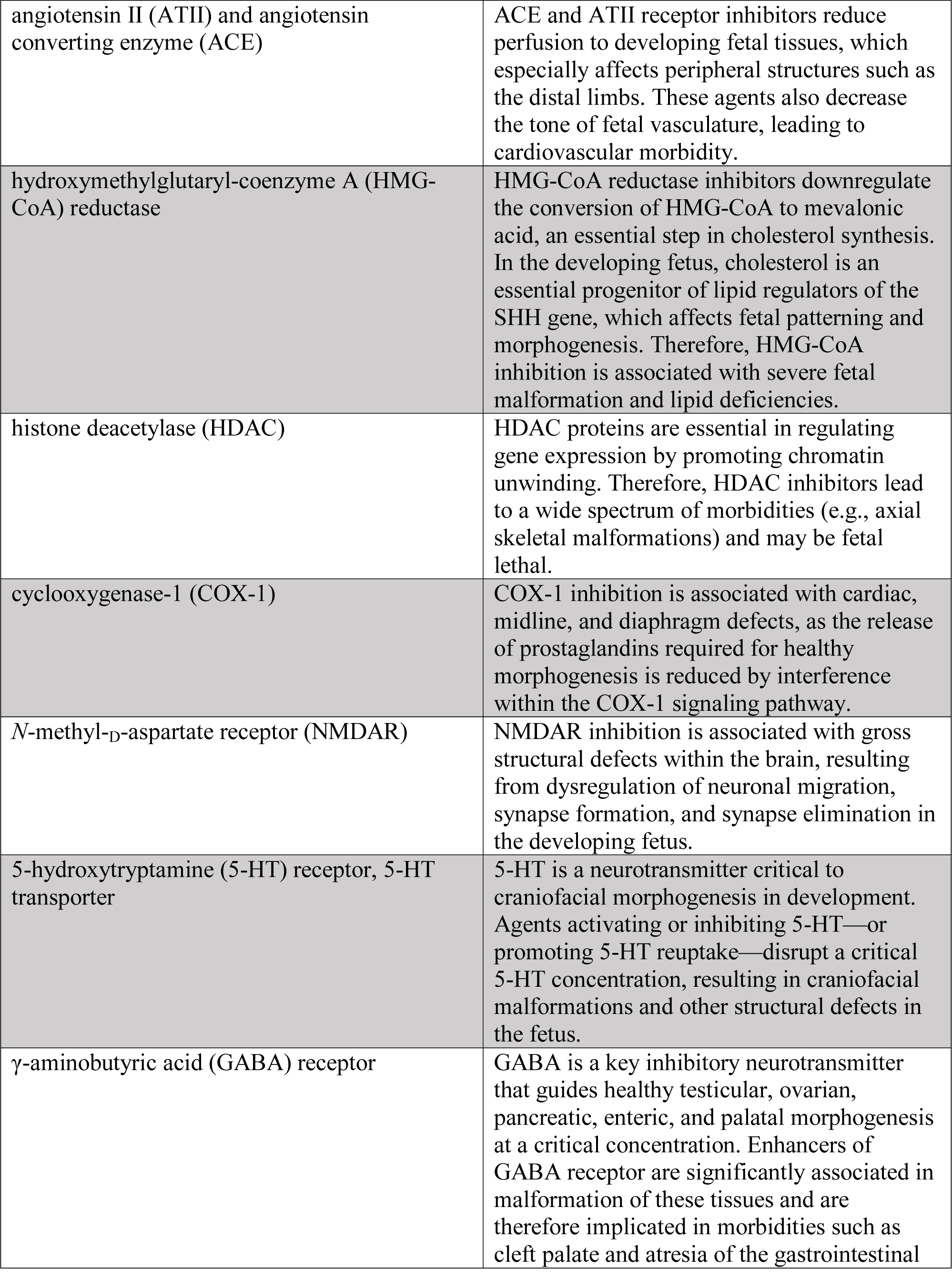
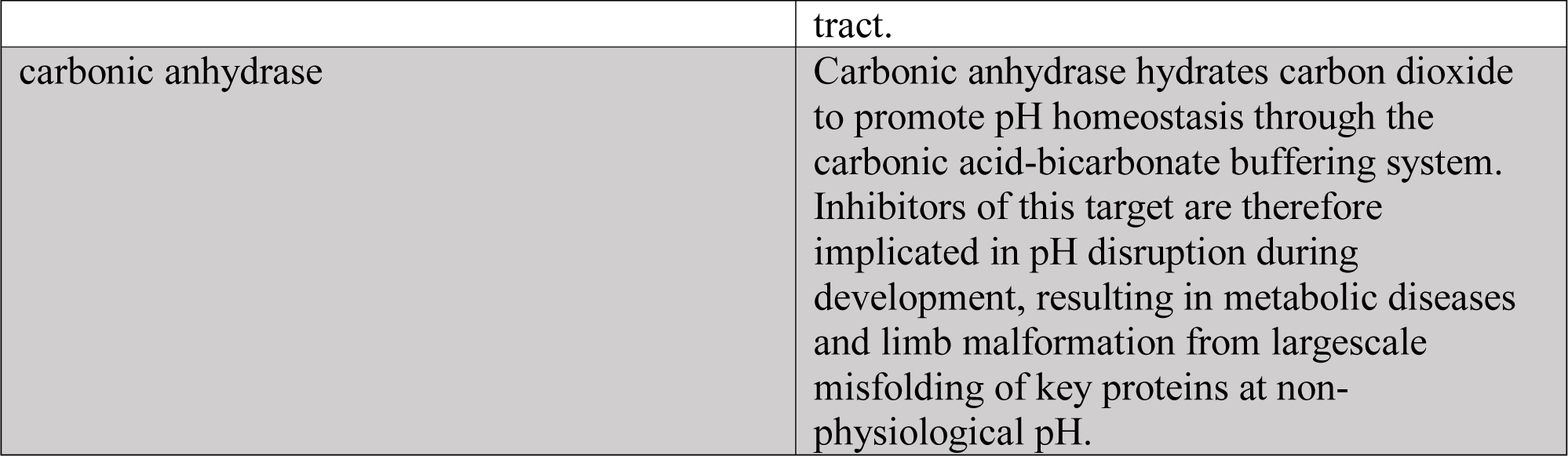
Teratogenesis converges on a limited subset of targets^19,44–53^.

### 1.3. Machine learning in maternal-fetal medicine

The inherent contradiction between the limited target rationale for teratogenesis and the extent of uncertainty that guides prescribing behavior for gravid populations speaks to the need for more rigorous predictions of small molecule teratogenicity. Furthermore, computational modeling on healthcare data is the most accurate method of predicting drug safety in pregnant women, given that phase I trials are unethical for expectant populations and animal models are inherently limited for studying human health^12, 13, 24^.

*Classification algorithms* are optimized to identify patterns between associated data sets (such as binding affinity and phenotype data for a cytotoxic target)^25–28^, suggesting that machine learning (ML) classifiers may play a pivotal role in systematically establishing relationships between maternal drug history and adverse fetal outcomes^29–31^. While these models are not intended as a replacement for existing physician knowledge of responsible prescriptive practice^32^, ML classifiers offer an attractive opportunity to discover meaningful relationships within existing biomedical data than could result in meaningful POC conclusions.

There have been few previous studies leveraging this brand of artificial intelligence (AI) for predicting iatrogenic fetal toxicity. Of these select investigations, a majority have focused solely on population-level, patient-derived data to discover adverse outcomes from maternal medication history and neonatal disease information^17,30,31,33,34^. In 2017, Boland *et al*. reported on a successful ML algorithm for parsing electronic health record (EHR) data to develop data-driven definitions of adverse drug outcomes associated with class C teratogenicity; the authors focused their modeling on congenital disease and fetal death phenotypes^35^. Studies with similarly-limited scope that analyzed insurance claims data are also available^30, 33, 36^. Recognizing the additional predictive power of chemical data for teratogenicity, Baker *et al*. published a ML model for the identification of compounds implicated in cleft palate formation from existing toxicology high-throughput screening (HTS) bioassay data and information on chemicals implicated in cleft palate phenotype identified from systematic literature review. This allowed the authors to identify biomarkers with high positive predictive value for cleft palate and further elucidate chemical exposure-adverse outcome clusters^17^.

In this study, we report on a previously-unattempted, unbiased (phenotype-agnostic and target-agnostic) approach to predicting teratogenicity by identifying chemical and biochemical factors that predispose a chemical to increased teratogenic risk. Given significant limitations in established teratogenicity scoring criteria, we propose a novel application of ML to develop a teratogenicity quantitative structure-activity relationship (QSAR)^37^. By leveraging drug structure, meta-structural elements like molecular energetics, and real-world bioactivity data, we attempt to predict the teratogenic risk of drugs potentially prescriptible in pregnancy.

## Materials and Methods

Our teratogenicity QSAR accesses chemical and bioassay data to predict a teratogenicity score for compounds that are prescriptible in pregnancy and to identify patterns within drug-specific information that predispose a drug towards an increased risk of fetal toxicity.

Broadly, we leverage three layers of drug data to accomplish these tasks:

1. The inherent structure of each drug, as encoded by several classes of chemical fingerprints^38^ that capture upwards of 1,024 structural features of each molecule
2. Meta-structural features for each drug, including druglikeness, predicted molecular energetics, and mutagenicty—as calculated from the Molecular Operating Environment (MOE)^39^, an industrial-grade chemical computing software—and mutagenicity data from predictive models of the Ames test^40^
3. Antagonist-mode toxicology assay data from Tripod, a public-facing collection of HTS data from the Toxicology in the 21st Century (Tox21) initiative of the National Institutes of Health^41,42^, on all targets listed in Table 2 with simultaneous coverage in Tox21

We employed version 3.5.3 of the R integrated development environment (https://www.r-project.org/)^43^ for parsing all chemical and bioassay data and implementing and tuning unsupervised and supervised ML models for associating these data.

### 2.1. Mining structure and teratogenicity data

DrugBank 5.1.0 (https://www.drugbank.ca/) is a publically-available drug encyclopedia developed by the University of Alberta. It contains comprehensive entries of more than two hundred (200) data fields for 9,099 small molecules of known structure that have passed phase I of an existing RCT. Each DrugBank entry contains structured information on compound structure, MOA, existing formulations, and drug marketing history, among other clinically-relevant datasets^44^. DrugBank is a self-described cheminformatics resource^45^; therefore, the pharmacopeia provides a highly pliable application programming interface (API), which allows for easy data mining and extraction. Given the comprehensiveness of the DrugBank database, as well as its amenability for data-driven analyses, we extracted the structures of all 9,099 DrugBank entries as three-dimensional spatial data files (3D-SDFs).

To obtain relevant, FDA-compliant teratology data, we interrogated SafeFetus (https://www.safefetus.com/)46, a registry for expectant mothers hosting the largest publically-available repository of structured, FDA-aligned teratogenicity scores. Therefore, we extracted teratogenicity scores for all 652 eligible drugs from SafeFetus.

Teratology data are not routinely published, and many large pharmacovigilance databanks like FDA’s DailyMed (https://dailymed.nlm.nih.gov/dailymed/)47, do not present all teratology information in structured fields—as required for computing on this information—and have inflexible APIs for data extraction.

### 2.2. Layer 1: Leveraging drug structure for predicting teratogenicity

From DrugBank 5.1.0, all 9,099 small molecule structure files were mined in SDF format. To ensure that DrugBank structure files were not corrupted in the extraction process, the SDF set was imported into the LigPrep graphical user interface of Schrödinger 2018-2 (https://www.schrodinger.com/)48, a suite of chemical computing software that enables predictive modeling in structure-guided pharmacological studies. Validating by visual inspection that all DrugBank files were chemically-valid, the SDF set was imported to R. Then, using the cheminformatics toolkits ChemmineR (*CRAN: ChemmineR*)^49^ and Rcdk (*CRAN: Rcdk*)^50^, the SDF set was converted to twelve (12) classes of chemical fingerprints, encodings of chemical structure as thousand-dimensional matrices that record the presence of absence of distinctive chemical motifs, including topological torsions, R/S stereochemistry, common functional groups, Brønsted-Lowey acidity/basicity, general acid/base catalysts, and other salient chemomarkers. This fingerprinting process is only valid for organic small molecules; therefore, all inorganic agents were automatically parsed from our drug set by the ChemmineR and Rcdk fingerprinting algorithms^38^. Thus, fingerprinting allowed us to access comprehensive, structured information on nearly nine thousand (9,000) small molecules and one-hot encode this information.

As noted above, we obtained FDA-compliant teratogenicity data from SafeFetus, the largest publically-available source of structured FDA teratogenicity scores with an API. Integrating the data sets for teratogenicity and drug structure in R, we obtained *N* = 611 drugs with information on both structure and teratogenicity.

We then developed multiple label classification strategies for teratogenicity scores, based on the nature of FDA teratology scores and a bibliostatistic search. One set of teratogenicity scores for all 611 drugs was aligned according to native FDA schema. A second set of scores was redefined as a three-pronged scale of bins: “Clinically Acceptable Risk” (scores A/B), “Moderate Risk” (score C), and “Clinically Unacceptable Risk” (scores D/X). A third scale was defined by a systematic literature search of the Embase medical library system (https://www.elsevier.com/solutions/embase-biomedical-research)^51^ and a Cochrane review (https://www.cochrane.org/evidence)52 for the term “teratogenic.” Visual inspection of nearly 16,000 resulting articles revealed strong association of the term “teratogenic” with a mention of FDA scores C, D, and X. Therefore, this binary scale was defined by “Non-Teratogenic” (scores A/B) and “Teratogenic” (scores C/D/X) classes.

#### 2.2.1. Unsupervised modeling

First, to discover clustering relationships between teratogenicity and drug structure, the Barnes-Hut implementation of the t-Distributed Stochastic Neighbor Embedding (t-SNE) procedure^53^ was enacted on all combinations of fingerprint and teratogenicity score data sets, including a combined, non-redundant, and feature-prioritized set of all chemical fingerprints. t-SNE is a dimensionality reduction procedure, which can plot all dimensions of drug structure against all dimensions of teratogenicity for all drugs included our data sets. The presence of tight clusters in a t-SNE plot indicates dependency between the plotted variables^54,55^.

Of the t-SNE combinations we attempted on our structure and teratology encodings, the t-SNE plot generated with 1,024-dimensional Morgan fingerprints^56^ and a binary classification of teratological risk showed the strongest clustering relationships. Clusters were identified by visual inspection, with each point within a cluster representing a drug. Hence, we mapped points within each discrete t-SNE cluster in reverse, from t-SNE space to its associated DrugBank entry.

Noting that all points within each cluster were consistent with a salient chemical functionality within the component drug structures, and that all cluster component drugs belonged to the same class, we considered our identification of meaningful clusters to be successful. Performing systematic literature review on each drug class identified as strongly associated to the presence or absence of elevated teratogenic risk, we noted that select drug class—teratogenicity score relationships identified by our model were verified in clinical decision-making tools like UpToDate (https://www.uptodate.com/contents/search)57 and Medscape (https://www.medscape.com/)58. However, several structure-teratology relationships identified by t-SNE appeared contentious in relevant literature: sufficient human data are not available to accurately classify the class of drugs distinguished by the t-SNE-identified chemical functionality as teratogenic or safe. We present a deeper discussion of the contribution of our t-SNE findings to these debates in the “Results and Discussion” section of this publication. The most consistent t-SNE plot and the functionalities it identified as significantly associated to the presence or absence of teratological risk are also shown in Figures 2 and 3 in the “Results and Discussion” section.

Noting that multiple structure-teratogenicity relationships resulting from our t-SNE analysis were validated in the literature, we considered our unsupervised ML model to be a successful proof-of-concept experiment. This—along with meaningful multiclass ROC analysis for structure-based predictions of teratogenicity (AUC = 0.8)—suggested that chemical structure has sufficient predictive power for a drug’s teratological risk to warrant the development of a supervised model for predicting teratogenicity from chemical fingerprints.

#### 2.2.2. Supervised modeling

Given that t-SNE successfully and consistently identified moieties that might predispose a drug towards an increased risk of teratogenicity, we decided to enable a supervised ML model that can prospectively predict a drug’s teratogenicity score from structural information. Using the R package Caret (*CRAN: Caret*)^59^, we developed three (3) models with inherent five (5)-fold cross validation (CV), such that we obtained test set accuracy on running each model. These models included Random Forest^60^, Extreme Gradient Boosting^61^, and Gradient Boosting Machine (GBM)^62^. Testing these models with five-pronged, FDA-adherent teratogenicity scores, we found that GBM yielded the highest predictive accuracy. Therefore, we re-trained our GBM model with the trivariate, clinically-oriented teratology scale described above and obtained higher accuracy for this model than for the GBM trained on five-dimensional labels. For all models, we optimized hyperparameters using a large grid search within Caret.

### 2.3. Layer 2: Curating meta-structural information for exploratory analysis

After deriving a successful model for predicting teratological risk from drug structure, we sought to increase the predictive accuracy of our GBM by supplementing our features with information on “meta-structure^63^.” These factors included the following variables, which were calculated for all 611 sampled drugs within MOE (https://www.chemcomp.com/Products.htm)39, a suite of industry-grade chemical computing software for computer-aided molecular design. Each of the following meta-structural sets was encoded by chemically-significant cutoffs when available (e.g., druglikeness benchmarks from Lipinski’s Rule of Five (RO5)^64^) or cutoffs determined from ROC analysis of extracted data):

Druglikeness: the adherence of each molecule to Lipinski’s Rule of Five restrictions on the number of hydrogen-bond acceptors, hydrogen-bond donors, octanol-water partition effects, total polar surface area, molecular weight, and number of rotatable bonds for an attractive drug candidate^64^
Energy of the Highest Occupied Molecular Orbital (HOMO): a quantum chemistry metric of the tendency of a molecule to donate an electron, as a proxy for drug stability and tendency to generate mutagenic free radicals^65^
Energy of the Lowest Unoccupied Molecular Orbital (LUMO): a quantum chemistry metric of the tendency of a molecule to accept an electron, as a proxy for drug stability and tendency to generate mutagenic free radicals^65^
Mutagenicity score, as calculated from in-built predictive models of the Ames test^40^
pKa and most basic pKa

#### 2.3.1. Unsupervised modeling

Combining MOE calculations for the above variables and all structural data sets, we performed feature selection within Caret to remove redundancy and highly-correlated features within the integrated descriptor set. Then, we re-executed t-SNE on binary teratogenicity scores, with the hope of identifying new clustering relationships between physiochemical features and teratogenicity.

#### 2.3.2. Supervised modeling

GBM with five-fold CV was re-executed with a three-pronged set of teratogenicity scores and feature-prioritized structural and meta-structural information. Hyperparameters were optimized by large grid search within Caret^59^.

### 2.4. Layer 3: Repurposing Tox21 HTS Data on Teratogenic Targets

Given that teratogenicity has well-identified target rationale, we decided to leverage existing, real-world bioassay information for all targets implicated in teratogenesis (as described Table 2) and previously screened through the Toxicology in the 21^st^ Century Initiative (Tox21) of the National Institutes of Health (https://ncats.nih.gov/tox21)41. Tox21 leverages HTS of millions of bioactive compounds—including most common pharmaceuticals—in thousand well-plate, cell-based assays. While this HTS platform is not teratogenicity-specific, it does contain information on targets implicated in teratogenesis^66^.

Scoping all information available on Tripod, the public-facing data browser of Tox21 (https://tripod.nih.gov/tox21)42, we extracted antagonist-mode RAR and HDAC data for supplementation of our model. RAR data were derived from murine embryo fibroblast cells (C3H10T1/2, American Type Culture Collection, Manassas, Va., USA), and HDAC data were obtained from human colorectal carcinoma cells (HCT-116, American Type Culture Collection, Manassas, Va., USA). Assay protocols are available from the Tripod website specified above.

Data available from Tripod include bioactivity for a given target (encoded as “inactive,” “active”), curve class, IC_50_, efficacy, and Hill coefficient. Of these variables, we studied curve class, IC_50_, and efficacy as proxies of binding affinity of each sampled compound for RAR and HDAC. Therefore, all compounds with available structure, teratogenicity score, and RAR/HDAC HTS coverage (*N* = 128) were probed by t-SNE and GBM. Data were one-hot encoded using standard bioactivity cutoffs for drug development (i.e., curve class ≠ 4, IC_50_ ≤ 20 µM, efficacy ≤ −50%)^67–69^.

#### 2.4.1. Unsupervised modeling

Combining MOE calculations for the above assay data and all structural and meta-structural data sets, we performed feature selection within Caret to remove redundancy and highly-correlated features within the integrated descriptor set. Then, we re-executed t-SNE on binary teratogenicity scores, with the hope of identifying new clustering relationships between assay data and teratogenicity.

#### 2.4.2. Supervised modeling

GBM with five-fold CV was re-executed with a three-pronged set of teratogenicity scores and feature-prioritized structural, meta-structural, and biochemical assay information. Hyperparameters were optimized by large grid search within Caret.

We are committed to open-source science. Code that we developed to execute this protocol is available through the following GitHub repository: https://github.com/apchalla/teratogenicity-qsar.

## Results and Discussion

In this manuscript, we present a first-in-kind application of ML to identify structural, meta-structural, and bioassay performance factors that predispose a drug towards increased teratogenic risk. We developed a model to prospectively score a drug’s teratogenicity from these drug-specific factors. Because our workflow is anchored in computing, our methods apply algorithmic rigor to studying teratogenicity, a contrast to many non-systematic studies which have historically dominated this space.

### 3.1. Summary of key results

#### 3.1.1. Unsupervised learning outcomes

We found that drug structure is a good predictor of teratogenicity, as multiclass ROC analysis between 1,024-dimensional Morgan fingerprints and a three-pronged teratogenicity metric gave AUC = 0.78 (Figure 1). This result validates our hypothesis that a “form-fits-function” argument is valid for predicting teratogenicity from homology between drug structure and pharmacophore biochemistry among targets implicated in teratogenesis.

**Figure 1:**
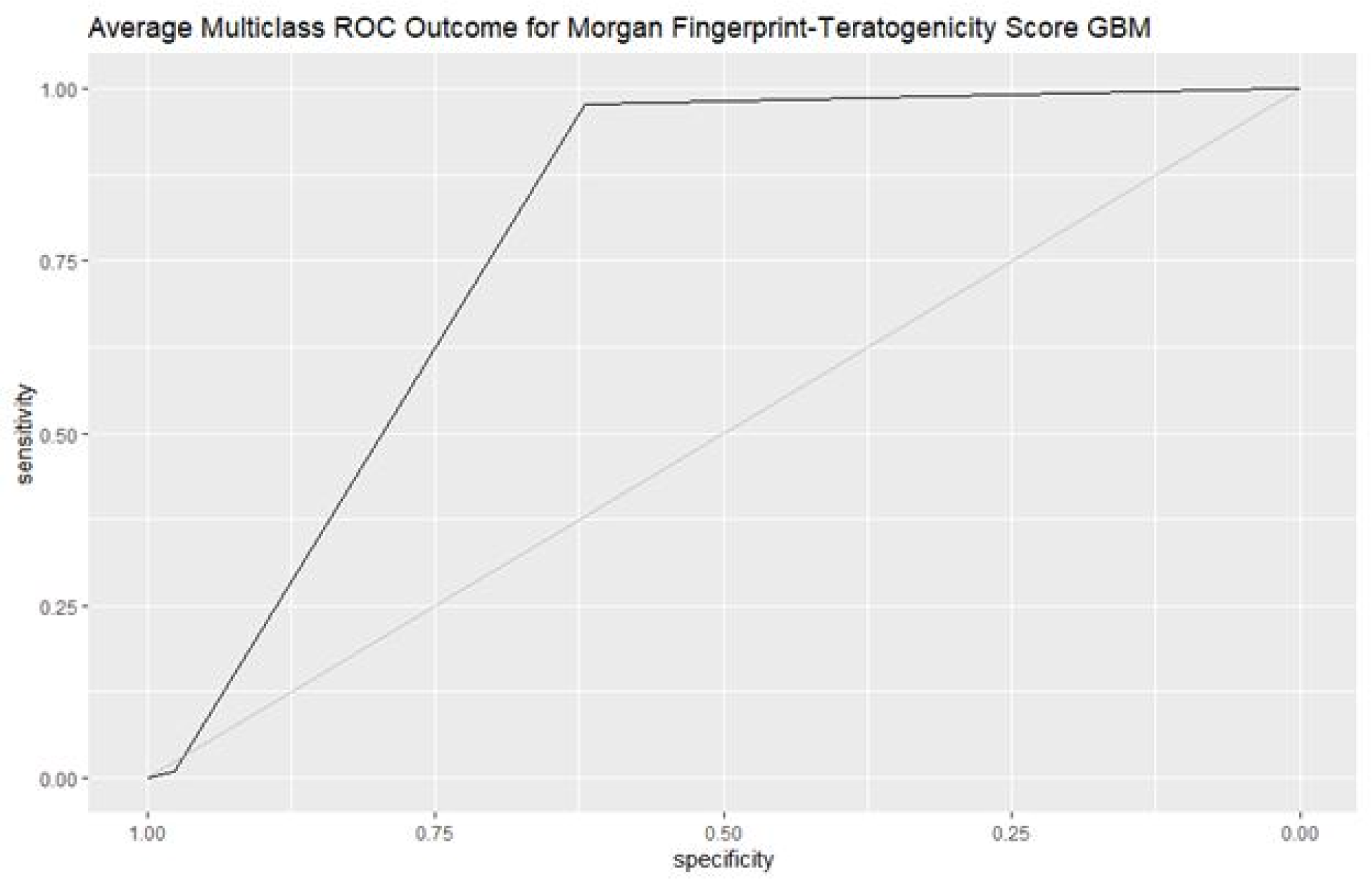
ROC analysis suggests that 1,024-bit Morgan fingerprints have good predictive accuracy for teratogenicity (AUC = 0.78). This plot was generated using the R package pROC (*CRAN: pROC*)^101^.

From t-SNE analysis between drug structure and a binary encoding of teratogenicity (Figure 2), we discovered clusters of teratogenic risk and the absence thereof, which are partially validated within existing clinical literature (Figure 3). Though t-SNE contains noise across most of the diminished structure-teratogenicity landscape, the clusters we identified by visual inspection were consistent in teratogenic risk. A reason for the limited tightness of the observed clustering behavior may involve dimensionality mismatch between structure and teratogenicity data sets, given that we plotted 1,024 structural motifs against only two (2) teratogenicity scores. However, since generating ∼10^3^ independent teratogenicity scores and reducing chemical structure to ∼10^1^ categories are both unfeasible (this would remove the clinical and chemical significance of the respective data sets), we cannot address this probable cause of loose clustering by adjusting the form of the data we seek to associate. Despite these issues, our t-SNE step was a successful proof-of-concept experiment, as we discovered functionalities that are known to be highly fetal toxic and those that are known to be safe through this procedure.

**Figure 2:**
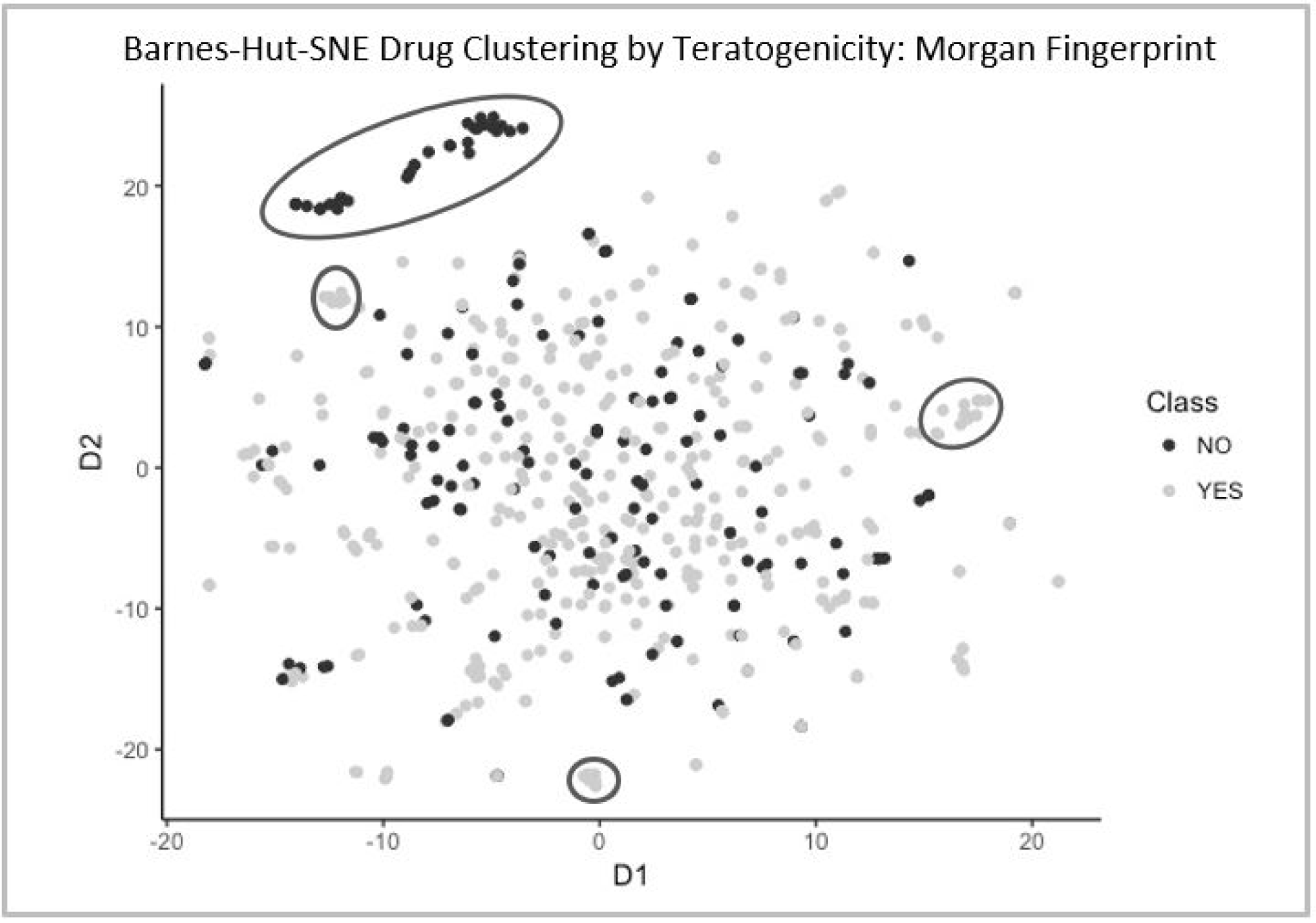
t-SNE—when enacted on a 1,024-bit representation of the Morgan class of chemical fingerprints and a binary classification of teratogenicity (“YES” (class A/B), “NO” (class C/D/X))—reveals small clusters that indicate potential structure-teratogenicity relationships. This plot was generated using the R package Rtsne (*CRAN: Rtsne*)^102^.

**Figure 3:**
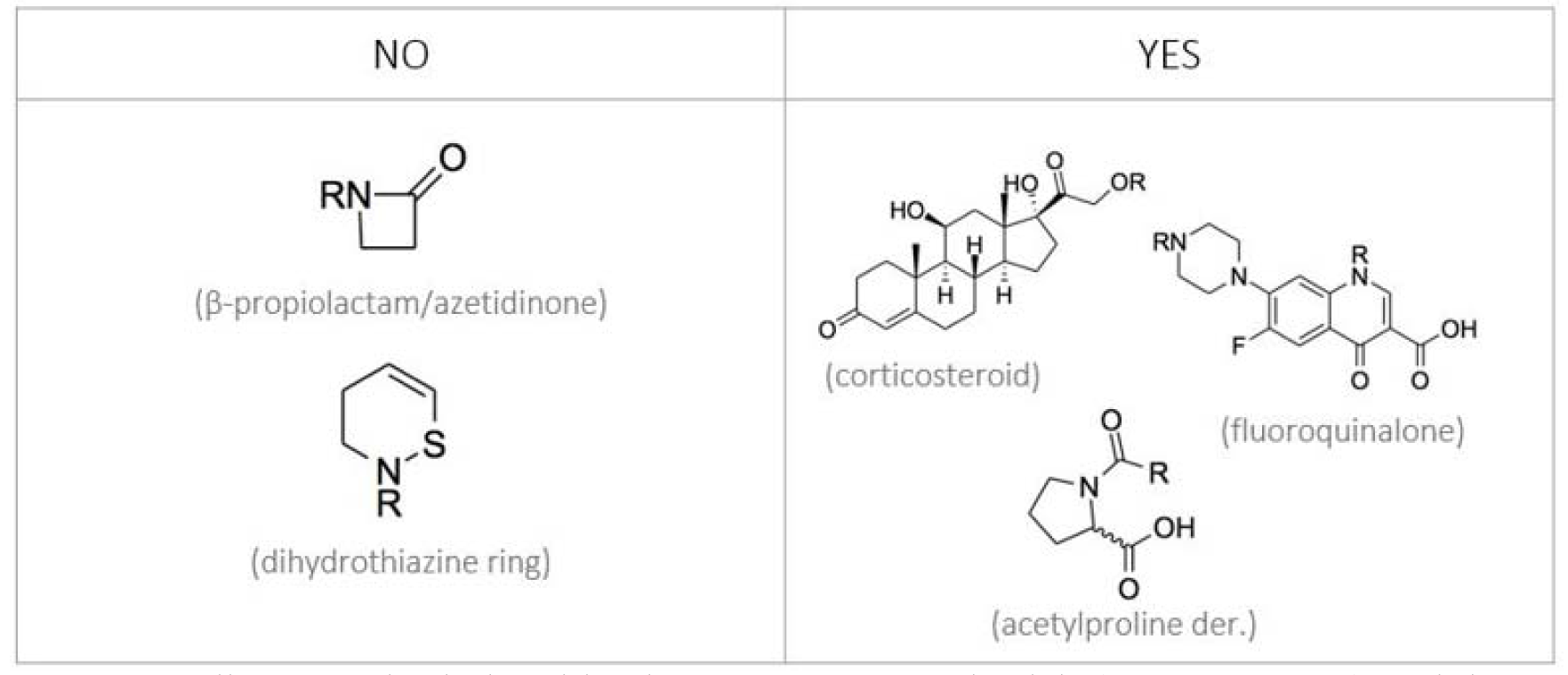
We discovered relationships between teratogenic risk (“YES”, “NO”) and the presence of distinct chemical functionalities from consistent structure-teratogenicity points within each discrete t-SNE cluster.

Beyond these validated associations, we also discovered new structure-teratogenicity relationships that might have application in clarifying cases of suspect toxicity risk in the clinical literature. Indeed, our analysis reveals five motifs that are distinctive among cohorts of molecules identified as “teratogenic” and “non-teratogenic.” Both moieties in the “NO” cluster are components of cephalosporins, which include a group of broad-range antibiotics known to be safe for pregnant mothers (class B)^70–73^. Two distinctive functionalities distinguish cephalosporins from other classes of drugs: the presence of an azetidinone group and a dihydrothiazine ring^74^. Therefore, as there features distinctively establish cephalosporin identity—which is non-teratogenic—it is reasonable to assert that the azetidinone functionality and dihydrothiazine ring are non-teratogenic chemomarkers in this case. We recognize that the burden of evidence is significant to claim that these motifs demonstrate protective effects. Instead, we suggest that our results warrant more involved analysis of these potentially protective moieties.

In contrast, similar analysis of “YES” clusters reveals three teratogenic chemomarkers, including corticosteroids, fluoroquinolones, and acetylproline derivatives. While fluoroquinolones are documented teratogens^75–78^, there is contention on the toxicity of steroid derivatives^79–81^, as well as prolinated compounds^82–84^. Our model adds to this discussion by arguing that steroid derivatives are indeed teratogenic.

We reasonably assume that the “YES” functionalities in Figure 3 are the source of teratogenicity within molecules that contain them, given that these moieties are distinctive. This conclusion requires MOA validation; however, as with fluoroquinolones, available phenotypic data appear to support our conclusions on functional group toxicity.

Drawing on these mappings also allows us to evaluate new trends in drug development; namely, we can extrapolate functional group mappings towards drug development targets in the anti-hypercholesterolemic space. Pregnant women with high cholesterol are not advised to take statins, as these drugs are antagonists of HMG-CoA reductase, restricting fatty acid synthesis in a developing fetus (Table 1)^19,85–87^. Statins contain a fluorobenzene motif, which our model predicts to be the core teratogenic functionality within these drugs. As of date, only one small-molecule anti-hypercholesterolemic drug, ezetimibe (Zetia), does not belong to the statin class of drugs^88^. Instead, ezetimibe contains a central azetidinone group and has been noted in reduced teratogenicity across the expectant population, as compared to statins^89^. Given that we identify azetidinone-containing drugs to carry potential protective effects, this result further edifies the results from our model and speaks to the applicability of structure-teratogenicity relationship modeling similar to that in this paper to inform downstream, data-driven inquiries into drug safety for expectant populations. We emphasize that expansion of this study and downstream mechanistic studies are required to fully substantiate our observations.

#### 3.1.2. Supervised learning outcomes

Our GBM predicts three classes of teratogenicity with 64.7% accuracy (SD = 3.0%) when trained on 1,024-dimensional Morgan fingerprints. Thus, our model achieves nearly double the predictive accuracy as a blind, probabilistic control for the same trivariate predictive task; QSAR accuracy enrichment is nearly 32% on these baseline predictions. Because there exist no other structure-activity relationships, meta-structure-activity relationships, or structure-assay-activity relationships published in this space, we assert our model as a first attempt at applying drug-inherent information towards predicting teratogenicity.

### 3.2. Ontological limitations and barriers to data-driven studies in pregnancy

While the results above appear promising, the data that we queried in this investigation present significant ontological challenges. These problems drastically reduce the sample size of all drug-specific teratology probes and present significant barriers to translational science, as we explain below.

In this study, we encountered problems with procuring teratogenicity information, given that teratology reference data are not published and updated in the relevant clinical literature very often. Furthermore, existing clinical decision-making tools like UpToDate and Medscape do not have APIs and contain contradictory teratology information that is not available in structured formats—as is required for systematic, retrospective data analysis and ML modeling. FDA resources containing teratology data are also not published in structured formats amenable for computational research, despite the availability an API for FDA pharmacopeias like DailyMed. For this investigation, the consequence of this limitation in available teratogenicity data was a significant reduction in drug sample size, as available to t-SNE and GBM. Though we used one of the arguably most powerful chemical computing software programs currently available (i.e., MOE), we encountered sparsity in meta-structural predictions within our limited subset of drugs with available structure and teratogenicity information. This restricted the power of our meta-structural t-SNE and GBM probes, resulting in no test power for a feature-selected meta-structural and structural feature set.

Despite the gravity of the inherent uncertainty within available teratogenicity scoring criteria and limited target rationale for teratogenesis, there exist no teratology-specific HTS platforms. Though large toxicology HTS programs like Tox21 have screened targets that overlap with those in Figure 1, this intersection remains small: only two (2) targets have coverage through Tox21. Therefore, though real-world bioactivity information is inherently powerful, we were able to access data on only two (2) relevant targets, and for only 128 drugs with structure and available teratogenicity data and assay information. Only sixty-four (64) drugs had information available for both RAR and HDAC, available structure data, and a known teratology score. Hence, a major reason why the addition of Tox21 HTS data did not improve predictive accuracy or t-SNE clustering over a purely structural model was limited sample size. This issue remains intractable, given the inherently limited data resources currently existing available and little action on the part of data providers to address these quality issues.

## Conclusions

Current standards of evaluating small molecule teratogenicity are inherently unsystematic and driven on a lack of human data. This informs irresponsible prescribing behavior at the POC, reducing the quality of care for pregnant women and their developing fetuses. However, given the rigor of rules-based ML classification algorithms and limited “on-target” rationale for teratogenesis, there is potential to systematically predict a compound’s risk for fetal toxicity by leveraging AI on drug-specific information, such as drug structure, meta-structure, and existing real-world bioassay data, as a proxy for binding affinity to teratogenic targets.

In our study, we assert that drug structure is a good predictor of teratogenicity, using ROC analysis, unsupervised ML (t-SNE), and a supervised GBM to discover relationships between chemical functionalities within drugs prescriptible in pregnancy and existing teratogenicity information. This allowed us to identify moieties that appear to predispose a drug towards an increased chance of teratogenicity, based on existing use cases that are salient in relevant clinical and drug development literature. We also identify significant barriers to translational research in this space as rationale for the limited utility of existing meta-structural and toxicology HTS platforms for teratogenicity prediction tasks. The importance of these ontological considerations cannot be overstated in considering future research to improve the quality of data-driven maternal-fetal medicine.

Our team of investigators has formed a first-in-kind research collaboration of engineers, informaticians, and clinicians dedicated to the development of computational tools to predict adverse drug outcomes in pregnancy from existing healthcare data on pregnant populations and *in vitro* drug exposure models that are more representative of pregnant human physiology than the *in vivo* animal platforms currently employed in this space. This group—called Modeling Adverse Drug Reactions in Embryos (MADRE)^13, 90, 91^—proposes refinement of the teratogenicity QSAR reported in this manuscript by harnessing a more continuous spectrum of relevant phenotype information. Given that data quality and availability issues with teratogenicity scores restricted the scope of this study, we propose a medication history-wide association study (MedWAS) that can leverage billing-encoded, population-level EHR data as a label set. The benefit of MedWAS over QSAR is increased flexibility: associative study model architecture would not necessitate classification of adverse outcomes into rigid bins, as the QSAR requires^37^. Therefore, MedWAS would not be restricted by the limited availability of FDA-encoded teratogenicity data, giving a larger sample size of drugs eligible for analysis and a more continuous spectrum of phenotype information through which to quantify teratogenicity. In turn, this allows for easier validation of associative outcomes *in silico* and *in vitro*, as compared to similar hits from QSAR. Indeed, drugs identified as teratogenic through MedWAS may be referred to our QSAR model for validation, and vice versa. We have begun work on this MedWAS and look forward to further exploring its intersections with our teratogenicity QSAR.

## Author Contributions

APC designed this study, performed all data extraction, ML analysis, and model tuning, and drafted this manuscript. ALB provided technical guidance on appropriate ML models for this study, assisted with model tuning, and helped interpret model results. MS helped design this study, provided access to Tox21 HTS data, provided technical guidance on model tuning, and helped interpret model results. TP provided technical guidance on the choice of features for this study, assisted with model tuning, and reviewed this manuscript. RRL provided technical guidance on the choice of features for this study, helped interpret model results, and reviewed this manuscript. ESL helped interpret model results and reviewed this manuscript. DMA helped interpret model results and reviewed this manuscript.

## Conflicts of Interest

We declare no competing interests relevant to the execution or outcomes of this study.

## Acknowledgements

We thank Asher Schachter, MD, Senior Vice President, Clinical, and Head of Pharmaceutical Sciences at CAMP4 Therapeutics, for sharing teratogenicity data that he extracted from SafeFetus.

Research reported in this publication was supported by the National Human Genome Research Institute of the National Institutes of Health under Award Number U54HG007963-05 and the National Center for Advancing Translational Sciences of the National Institutes of Health under Clinical and Translational Science Award Number U54TR02243-02. The content is solely the responsibility of the authors and does not represent the official views of the National Institutes of Health.

## Abbreviations

RCT: randomized controlled trial
FDA: United States Food and Drug Administration
POC: point of care
MOA: mechanism of action
ROS: reactive oxygen species
ML: machine learning
AI: artificial intelligence
EHR: electronic health record
HTS: high-throughput screening
QSAR: quantitative structure-activity relationship
MOE: Molecular Operating Environment
Tox21: Toxicology in the 21^st^ Century Initiative
API: application programming interface
3D-SDF: three-dimensional spatial data file
t-SNE: t-Distributed Stochastic Neighbor Embedding
GBM: gradient boosting machine
CV: cross-validation
RO5: Lipinski’s Rule of Five
HOMO: highest occupied molecular orbital
LUMO: lowest unoccupied molecular orbital
MADRE: Modeling Adverse Drug Reactions in Embryos
MedWAS: medication history-wide association study

## Notes

https://github.com/apchalla/teratogenicity-qsar.

